# Alpha-synuclein aggregation and dopaminergic neuron death in a new mouse model of Parkinson’s disease expressing human full-length and C-terminally truncated 1- 120 alpha-synuclein

**DOI:** 10.1101/2025.04.13.647581

**Authors:** F Longhena, Z Zheng, G Faustini, M Wegrzynowicz, E Carlson, N Casadei, O Riess, A Bellucci, MG Spillantini

## Abstract

Parkinson’s disease (PD) is neuropathologically characterized by the presence of Lewy bodies and Lewy neurites made of aggregated alpha-synuclein (aSyn). Several studies have shown that Lewy bodies contain C-terminally truncated aSyn which is more prone to fibrillation and enhances the aggregation of human full-length aSyn *in vitro*. Nevertheless, *in vivo* studies addressing whether and how human C-terminally truncated aSyn may exacerbate the pathological aggregation and toxicity of human full-length aSyn are currently lacking. In the present study we aimed to determine whether the co-expression of human full-length and 1-120 truncated aSyn would enhance human full-length aSyn aggregation and toxicity. To this end we generated and characterized two novel alpha-synuclein mouse models. Specifically, we first produced a mouse line expressing full-length human aSyn under its own promoter (BACo mice) in the absence of endogenous mouse aSyn. Then, the BACo mice were crossed with the previously described MI2 mice, which express human 1-120 truncated aSyn under the control of the tyrosine hydroxylase (TH) promoter on a mouse aSyn null backgroundin order to obtain the MI2BACo mice, which express both human 1-120 truncated and full-length aSyn in a mouse aSyn null background. We found that aSyn aggregation was present and increased with age in both mouse lines but was significantly higher in the MI2BACo mice. These mice exhibited progressive nigrostriatal accumulation of aSyn aggregates which were often composed of full-length and 1-120haSyn, the latter being specifically recognized by using a novel antibody raised against the 1-120haSyn. In addition, we observed that the presence of both truncated and full-length aSyn in MI2BACo mice led to a faster and more pronounced aSyn aggregation which associated with more severe dopaminergic striatal fiber degeneration and nigral neuron loss. Collectively, our results indicate that the expression of human C-terminally truncated aSyn exacerbates human full-length aSyn pathology and support that the novel MI2BACo transgenic mouse line represents an innovative experimental model to study the biological basis of PD and test novel therapeutic approaches.

## INTRODUCTION

Parkinson’s disease (PD), the most common movement disorder, is characterized by the loss of dopaminergic neurons in the nigrostriatal system, which is associated with the onset of motor symptoms (Hornykiewicz 1998). Neuropathologically PD is characterized by the presence of Lewy bodies (LB) and Lewy neurites (LN), proteinaceous inclusions containing filamentous alpha-synuclein (aSyn) aggregates forming in neuronal cell bodies and neurites, respectively (Spillantini et al. 1997, Spillantini et al. 1998). aSyn is a short protein of 140 amino acids, very abundant in the nervous system where it is mainly localized in pre-synaptic terminals and is involved in vesicle transport and neurotransmitter release (Sharma et al. 2023) through interactions with synaptic proteins such as VAMP2 and synapsin III (Syn III) (Zaltieri et al. 2015; Somayaji et al. 2020; Faustini et al. 2022). In line with its enrichment at synaptic terminals, several studies support that aSyn pathological aggregation is likely to initiate at the synapse, causing alterations in neurotransmitter release, progressing then to the cell body in a retrograde manner and ultimately leading to dopaminergic neuron death and motor symptoms (Kramer et al. 2007, Garcia-Reitbock et al. 2010, Schulz-Schaeffer 2010, Bellucci et al. 2016, Andica et al. 2018, Tozzi et al. 2021). Recent post-mortem studies have also shown that in PD synaptic loss is linked to axonal damage and severe aSyn burden (Frigerio et al. 2024).

We also found that expression of C-terminally truncated aSyn under the rat tyrosine hydroxylase (TH) promoter in aSyn null mice produced a retrograde degeneration of nigral neurons, reminiscent of the putaminal deafferentation observed by tractography and magnetic resonance imaging in PD patients (Garcia-Reitbock et al. 2010; Faustini et al. 2022; Andica et al. 2018; Lopez-Aquirre et al., 2023). C-terminally truncated aSyn is present in extracts from PD brains (Baba et al. 1998, Liu et al. 2005, Tofaris et al. 2005, Dufty et al. 2007, Prasad et al., 2012; Muntane et al. 2012; Moors et al. 2021) and it has been hypothesised that it could trigger full-length protein aggregation and LB formation (Kuusisto et al. 2003). Accordingly, several *in vitro* studies have shown that C-terminal truncation of aSyn promotes its faster aggregation (Crowther et al. 1998, Prasad et al. 2012, Bassil et al. 2016, Sorrentino et al. 2018, van der Wateren et al. 2018, Kumari et al. 2021, Zhang et al. 2022) possibly because its C-terminus is important for maintaining its unfolded state (Kumari et al. 2021). For this propensity to aggregate, of C-terminally truncated aSyn has been used to accelerate aSyn aggregation in transgenic mice (Tofaris et al. 2006, Daher et al. 2009, Hall et al. 2015; Wegrzynowicz et al. 2019). Another factor affecting human aSyn aggregation in transgenic mice is the presence of endogenous mouse aSyn that has shown to inhibit human aSyn aggregation (Fares et al. 2015; Luk et al. 2016, Ohgita et al. 2023). Probably this could be related to the fact that mouse aSyn fibrillar aggregates do not reproduce the structure of human filaments (Yang et al. 2022; Sokratian et al., 2024).

This suggests that the generation of aSyn transgenic mice expressing human aSyn in the absence of mouse aSyn can enable a better understanding of aSyn pathophysiology. We have previously reported two transgenic mouse models expressing 1-120 C-terminally truncated aSyn (1-120haSyn) under the control of the rat TH promoter in a mouse aSyn null background (Tofaris et al. 2006; Wegrzynowicz et al. 2019). The transgenic mouse line named MI2, shows progressive aSyn aggregation as well as nigrostriatal dopaminergic neuron dysfunction and loss. In the present study we describe a new mouse line, named BACo, expressing human full-length aSyn under its own promoter (Yamakado et al. 2012, Nuber et al. 2013) in a mouse aSyn null background (Specht et al. 2001). In order to determine whether the aSyn pathology was increased by the co-expression of human full-length aSyn and C-terminally truncated aSyn, we crossed the MI2 mice and BACo mice to generate the MI2BACo mouse line which expresses both 1-120 truncated and full-length human aSyn. Moreover, we generated a new polyclonal antibody (L91) that specifically recognizes 1-120haSyn and we used it to investigate its distribution and its interaction with full-length human aSyn. In the MI2BACo mice, we evaluated whether the co-expression of C-terminally truncated and full-length aSyn enhanced the degeneration of nigrostriatal neurons. We found that the presence of 1-120haSyn increases human full-length aSyn aggregation in the MI2BACo mice compared to BACo and MI2 mice. The L91 immunostaining often co-localized with Thioflavin S-positive (Thio-S) aggregates supporting that 1-120haSyn forms fibrillar aggregates. Furthermore, the MI2BACo mice show enhanced degeneration of nigrostriatal neurons at 12 months of age when compared to BACo mice. The results in the MI2BACo mice indicate that also *in vivo* 1-120haSyn leads to a faster aggregation of full-length human aSyn enhancing the pathological phenotype.

## MATERIAL AND METHODS

### Transgenic mice

aSyn BACo mice were generated using a *Bacterial Artificial Chromosome* (BAC) construct comprising a fused PAC AF163864 and BAC AC097478 containing the entire human *SNCA* gene locus with 28 kb 5′ - and 50kb 3′-flanking regions. This DNA construct was previously used to generate BAC transgenic mice (Yamakado et al. 2012) in a C57Bl6J background and aSyn BAC rats (Nuber et al. 2013). The same DNA construct was injected in C57BL6N mice to generate another BAC transgenic mouse (Wassouf et al. 2018). Using sperm from the BAC/C57Bl6N male mice we fertilised eggs from C57Bl6OlaHsd (OlaHsd) female mice generating a new BAC mouse in a OlaHsd background (BACo) without the endogenous mouse aSyn. Following breeding of the BACo mice to homozygosity we crossed them with MI2 transgenic mice to generate the homozygous MI2BACo line expressing both human 1- 120haSyn and human full-length aSyn in an endogenous mouse aSyn null background. Use of animals was performed under the Animals (Scientific Procedures) Act 1986, Amendment Regulations 2012, following an ethical review by the University of Cambridge Animal Welfare and Ethical Review Body (AWERB). PPL n PP4466737

### Human tissues

Five µm thick paraffin sections from the *substantia nigra* and *caudate putamen* from two patients with PD and two age-matched control subjects were obtained from the Parkinson’s UK Brain Bank (Imperial College London) and used for immunostainings. Handling of human tissue was according to the U.K. Human Tissue Act 2006 and is covered by the Cambridge Local Research Ethics Committee.

### Antibodies

The information about primary antibodies and dilutions used for the experiments described are listed in Table 1. L91 rabbit polyclonal antiserum was developed by Biosynth Laboratories Ltd, formerly Cambridge Research Biochemicals, by using a specific aSyn antigen that mimics the 1-120 C-terminally truncated protein.

| PRIMARY ANTIBODY | MANUFACTURER | WORKING CONCENTRATION |  |
| --- | --- | --- | --- |
|  |  | WB | IHC/IF |
| aSyn 4B12 (Human) | Thermo Fisher cat. MA1-90346 | 1:2000 |  |
| aSyn LB509 (Human) | AbCam cat. ab27766 |  | 1:500 |
| aSyn Syn1 (Total) | BD Traduction Lab cat. 610787 | 1:2000 | 1:600 |
| aSyn L91 (1-120 aa) | Biosynth Laboratories Ltd formerly Cambridge Research Biochemicals | 1:1000 | 1:500 |
| B-Actin | AbCam cat. ab8227 | 1:5000 |  |
| SNAP25 | BioLegend cat. 836304 |  | 1:500 |
| Synapsin III | SYSY cat. 106 303 |  | 1:500 |
| Tyrosine Hydroxylase (TH) | Merck Millipore cat. AB152 |  | 1:600 |

### Western blotting

Animals were euthanized via cervical dislocation. Brains were snap frozen in dry ice and stored at −80 °C. Brain regions (cortex, striatum, *substantia nigra*) were isolated, homogenized in 10% w/v RIPA buffer (50 mM Tris-HCl pH 7.5 + 1% NP-40 + 0.5% Sodium deoxycholate + 0.1% SDS + 150 mM NaCl + 5 mM EDTA, supplemented with phosphatase and protease inhibitors), and centrifuged at 4 °C at 14,000×g for 5 minutes. The supernatant was collected and the protein concentrations was determined using a BCA protein assay kit (Merck Millipore). After heating the samples at 95 °C for 5’ in Loading Buffer 4x (Thermo Fisher) supplemented with β-mercaptoethanol, proteins were separated by 4-12% SDS-PAGE gels (Bio-Rad) and transferred onto polyvinylidene difluoride (PVDF) 0.2 µm membranes (Merck Millipore). To block non-specific binding, the membranes were incubated with blocking solution (5% BSA in TBS with 0.5% Tween 20 (TBST)) for 1 hour at room temperature (RT). They were then incubated overnight at 4 °C with primary antibodies diluted in blocking solution. The following day, after three washes with TBST, the membranes were treated with peroxidase-conjugated secondary antibodies diluted in TBST (1:2000-4000, Dako) for 2 hours at RT, and the blots were visualized using a Chemi Doc MP imager (Bio-Rad) with West Dura Extended Duration Chemiluminescent Substrate (Thermo Fisher). Image Lab 5.1 (Bio-Rad Laboratories) was used for blot analysis.

### Immunohistochemistry

Mice were anesthetized via intraperitoneal injection of pentobarbital (Merial Animal Health) and then transcardially perfused with ice-cold PBS, followed by 4% PFA in PBS. The brains were post-fixed overnight in 4% PFA, then transferred to 30% sucrose in PBS with 0.1% NaN_3_ and stored at 4 °C. The brains were then frozen, and 30 μm sections were cut using a freezing microtome (Bright Instruments). Human or mouse sections were mounted onto glass slides, washed with PBS + 0.3% Triton X-100 (PBST) and antigen retrieval was performed in 10 mM sodium citrate buffer (pH 8.5) at 95 °C for 20 minutes. Endogenous peroxidase activity was inhibited by incubating the sections with 5% H_2_O_2_ + 20% methanol in PBST for 20’ at RT. Sections were then incubated with blocking solution (5% BSA in PBST) for 1 hour at RT, to reduce non-specific background and then with primary antibodies diluted in blocking solution overnight at 4°C. The following day, sections were incubated with biotinylated secondary antibodies (IgG Antibody (H+L) from Vector Laboratories) diluted in PBST (1:1000) for 1 hour at RT and peroxidase-based staining was developed using the Vectastain Elite ABC HRP Kit (45’ incubation at RT) and the DAB Peroxidase Substrate Kit (Vector Laboratories). Sections were then dehydrated, cleared in xylene, and coverslipped using DPX mounting medium. Sections stained with SG substrate were incubated with ready-made ImmPRESS^®^ HRP IgG PLUS Polymer Kit for 1 hour at RT, and ImmPACT^®^ SG Substrate Kit (Vector Laboratories). The sections were then dehydrated, cleared in xylene, and coverslipped as mentioned above. Pictures were taken using Olympus BX 53 light microscope or Zeiss Axiovert S100 and camera PCO Sensicam and analyzed with Fiji (NIH).

### Quantification of nigral TH-positive (TH^+^) neurons in Substantia Nigra

The number of TH^+^ cells was estimated using double-blind cell counting under bright-field microscopy with the optical fractionator method (Kirik et al. 2002, Faustini et al. 2018). Neurons from the substantia nigra pars compacta SNpc were analyzed with an inverted Zeiss Axiovert S100 microscope connected to a PC running the StagePro module of Image- ProTM Plus software (version 6.2, Media Cybernetics, Inc.), following the protocol described by Kirik et al. (2002). The entire mesencephalon was sectioned, with three 30 μm thick sections examined every 150 μm along the rostro-caudal axis. TH^+^ cells lateral to the medial terminal nucleus of the accessory optic tract, marking the medial boundary of the SNpc, were counted. The ventral tegmental area was excluded from these cell counts.

### Culture of SK-N-SH neuroblastoma cells overexpressing aSyn tagged with red fluorescent protein (RFP)

Human neuroblastoma SK-N-SH cells stably overexpressing full-length aSyn tagged with RFP (RFP-aSyn 140) or 1-120haSyn tagged with RFP (RFP-aSyn 120) were grown in Dulbecco’s modified Eagle’s medium (DMEM) with 1000 mg glucose/l (Sigma Aldrich) supplemented with 10% heat-inactivated fetal bovine serum, 100 μg/ml penicillin, 100 μg/ml streptomycin and 0.01 μM non-essential amino acids (Sigma-Aldrich). Cells were maintained at 37 °C under a humidified atmosphere of 5% CO_2_ and 95% O_2_.

For immunofluorescence, cells were plated on 24 wells plate (10.000 cells/well) and maintained in growth medium for three days, fixed for 10 min with 4% PFA in PBS, and stored at 4°C in PBS with 0.1% NaN_3_.

### Immunofluorescence

After perfusion and sectioning, 30 µm mouse brain sections were mounted on glass slides. Both mouse sections and fixed cells were permeabilized with 20% methanol in PBS for 20 min at RT, incubated for 1 hour at RT in blocking solution (5% BSA in PBST) and then with the primary antibody diluted in blocking solution overnight at 4 °C. The following day, samples were washed with PBST and incubated with the fluorochrome-conjugated secondary antibody (AlexaFluor 488-conjugated, Thermo Fisher) diluted in PBTS (1:1000) for 1 hour at RT. After three washes with PBST, slides were incubated with the second primary antibody for 2 hours at RT and then with suitable fluorochrome-conjugated secondary antibody (AlexaFluor 594-conjugated for mouse sections and AlexaFluor 647- conjugated for SK-N-SH cells, Thermo Fisher) diluted in PBST (1:1000) for 1 hour at RT. Cell nuclei were counterstained with Hoechst 33258 (Sigma-Aldrich) and then brain slices were incubated for 1 min at RT with TrueVIEW^®^ Quenching Kit (Vector Laboratories) to reduce autofluorescence non-specific signal. Sections were coverslipped with Vectashield^®^ mounting medium (Vector Laboratories) and analyzed by confocal microscopy (Zeiss LSM 900 Confocal Microscope).

### Thioflavin-S (Thio-S) staining

To visualize β-sheet-enriched aSyn aggregates, mouse brain sections were incubated with 0.05% Thio-S (Sigma-Aldrich) dissolved in 50% ethanol for 8 min at RT, followed by three washes with 80% ethanol to remove non-specific dye residues in the tissue. The sections background was then blocked for subsequent immunostaining with an antibody against aSyn and counterstained with Hoechst 33258, mounted on slides and analyzed using confocal microscopy (Zeiss LSM 900 Confocal Microscope).

### Confocal microscopy

Slides were imaged using LSM 900 Zeiss confocal laser microscope (Carl Zeiss) with the following laser sets: λ = 488/543/405 for double immunofluorescence and 543/430/405 for Thio-S staining. The height of sections scanning was 1 μm. Images (512 × 512 pixels) were then reconstructed using Zeiss ZEN Imaging Software (Carl Zeiss).

### Statistical analysis

Statistical differences between groups in positive particles size or positive area estimated by brain section staining or protein levels detected by western blot were assessed by using Student’s t-test or one-way ANOVA followed by Bonferroni’s multiple comparisons test (in case of >2 groups, n = 3-4 animals for each group for staining and western blot analysis). All data are presented as mean ± standard error of the mean (SEM) and statistical significance was established at P < 0.05.

## RESULTS

### The L91 antibody specific for 1-120haSyn efficiently discriminates between 1- 120haSyn and full-length human aSyn

In order to distinguish the truncated human 1-120 aSyn from the full-length human aSyn, both expressed in the MI2BACo mice, we raised an antibody that specifically recognizes the 1-120 truncation in aSyn. The antibody was generated using a peptide mimicking the 1-120 truncation in aSyn and its specificity was determined by immunofluorescence, immunoblotting and immunohistochemistry.

The specificity of the antibody was evaluated on the human neuroblastoma cell line SK-N-SH that overexpressed either full-length or 1-120haSyn tagged with RFP (SK RFP-aSyn 140 and SK RFP-aSyn 120). Fixed cell cultures were double stained with L91 and Syn1, the immunolabelling confirmed that L91 recognizes only SK RFP-aSyn 120 (Suppl. Figure 1G, L, J, O), while SK RFP-aSyn 140 was not recognized by L91 (Suppl. Figure 1V, AA, Y, AD). RFP-aSyn 140 cells were stained only by Syn1 (Suppl. Figure 1W, AB) that recognized also truncated 1-120haSyn in SK RFP-aSyn 120 cells (Suppl. Figure 1H, M). Blank consisted of cells incubated only with the secondary antibodies (Suppl. Figure 1A-E, P-T).

L91 was then tested on striatum and *substantia nigra* of 12month-old MI2BACo, MI2, BACo, and OlaHsd. While OlaHsd tissues did not show any specific signal (Figure 1G, H), BACo mice showed some background staining in the striatum (Figure 1F), where we cannot exclude the presence of truncated protein due to degradation of the full-length aSyn as previously reported in BAC rats using the same transgene (Nuber et al. 2013). On the contrary, tissue sections from MI2 and MI2BACo showed a strong immunopositive signal, with staining of neurons in the *substantia nigra* (Figure 1A,C) and displayed a puncta-like distribution indicative of synaptic staining in the grey matter in the striatum (Figure 1B, D). We then tested the L91 antibody on post-mortem brain of patients affected by PD and age matched controls. L91 stained numerous LBs (Suppl. Figure 2E,F) and LNs (Suppl. Figure 2C, D) in the substantia nigra of PD patients supporting the presence of 1-120 aSyn in these structures. In control brainwhere neuromelanin-positive grains could be seen, no staining for L91 was present. (Suppl. Figure 2A, B). To further demonstrate the specificity of the L91 antibody we performed western blot analysis of protein extracts from substantia nigra of 12 month-old BACo, 1-120haSyn, OlaHsd and C57Bl6J mice with endogenous aSyn expression (Suppl Figure 2G). Recombinant full-length and 1-120haSyn were loaded as positive controls. The L91 antibody recognized the 1-120haSyn band in the 1-120haSyn mice as well as the 1-120 recombinant aSyn but did not recognize the extracts from BACo, wildtype C57Bl6J mice or recombinant full-length aSyn.

**Figure 1.**
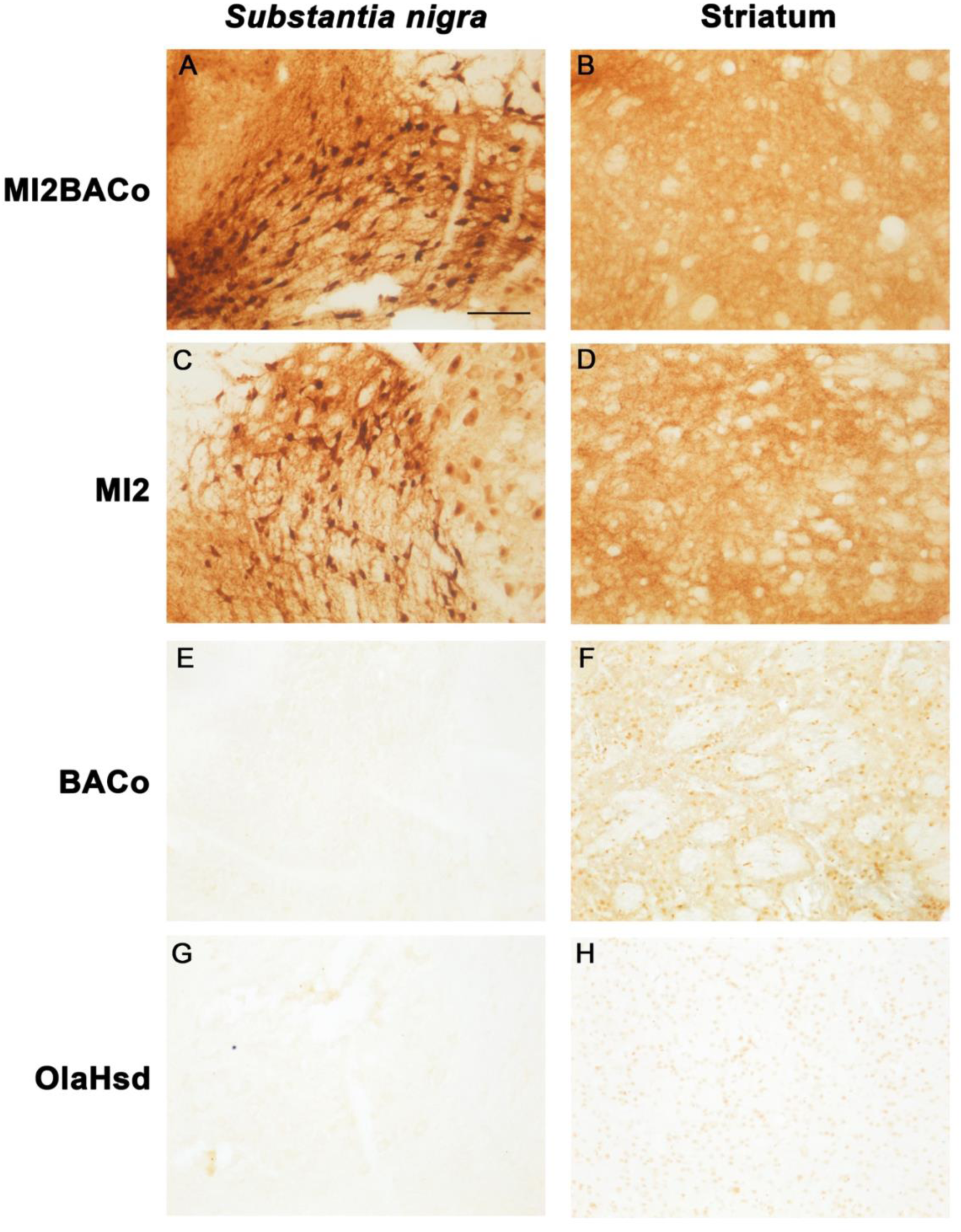
Characterization of a novel antibody (L91) specific for 1-120 truncated aSyn-(A-H) Representative images showing immunolabeling with L91, an antibody that specifically recognize 1-120 truncated and not full-length human aSyn, in the *substantia nigra* (A, C, E, G) and striatum (B, D, F, H) of 12 months old MI2BACo MI2, BACo and OlaHsd, mice. C57Bl6/OlaHsd do not express endogenous asyn and show no positive signal (G-H), while BACo mice show a faint signal in the striatum (F). MI2 (C, D) and MI2BACo (A, B) brains show staining for L91 and the presence of truncated aSyn specific signal in both the *substantia nigra* and striatum, as these transgenic mice both express human truncated 1-120haSyn under the control of rat TH promoter. Scale bar: 100 μm.

We then performed double immunofluorescence on the striatum of 12 month-old mice, using L91 and Syn1 antibody, that recognizes both full-length and truncated human aSyn (Suppl. Figure 3). We found that only MI2 (Suppl. Figure 3IE-H) exhibited co-localization between L91 and Syn1 antibodies. While BACo mice where positive only for the Syn1 antibody (Suppl. Figure 3A-D) further supporting the specificity of the L91 antibody for human 120haSyn. Interestingly, in the MI2BACo mice (Suppl. Figure 3I-L) Syn1 immunolabelling in was more extensive compared to L91 staining. The relative extent of co-localization between L91 and Syn1 in the MI2BACo mice appeared to be reduced, supporting that only a fraction of C-terminally truncated and full-length aSyn co-localize in these mice.

### Appearance of aSyn-positive clumps in the striatum of MI2BACo occurs earlier than in the MI2 mice and involves full-length human aSyn

MI2BACo mice express 120haSyn under the control of the rat TH promoter (Wegrzynowicz et al. 2019) on a BACo mouse background expressing human full-length aSyn without endogenous mouse aSyn. Western blot studies showed that the BACo mice, in line with what has been reported for other BAC aSyn models (Yamakado et al. 2012, Janezic et al. 2013, Nuber et al. 2013, Wassouf et al. 2018, Uemura et al. 2020) express about 2-fold higher amount of full-length human aSyn compared to the endogenous mouse aSyn in C57Bl6J wild type mice (Suppl. Fig 4). In order to determine whether the co-expression of 120haSyn and full-length aSyn enhances the formation of aggregates, striatal sections from 3, 6 and 12 months-old MI2BACo and BACo mice where immunolabelled with Syn1 antibody (Figure 2). We found that the size of the aSyn-positive clumps in the MI2-BACo mice was significantly increased when compared to BACo and MI2 mice already at 3 months of age (Figure 2J-L). At 12 months of age the size of aSyn clumps in the MI2 was similar to that of the MI2BACo mice at 3 months of age (Figure 2J,L), suggesting that full-length aSyn is sequestered and accelerates the formation of the clumps.

**Figure 2.**
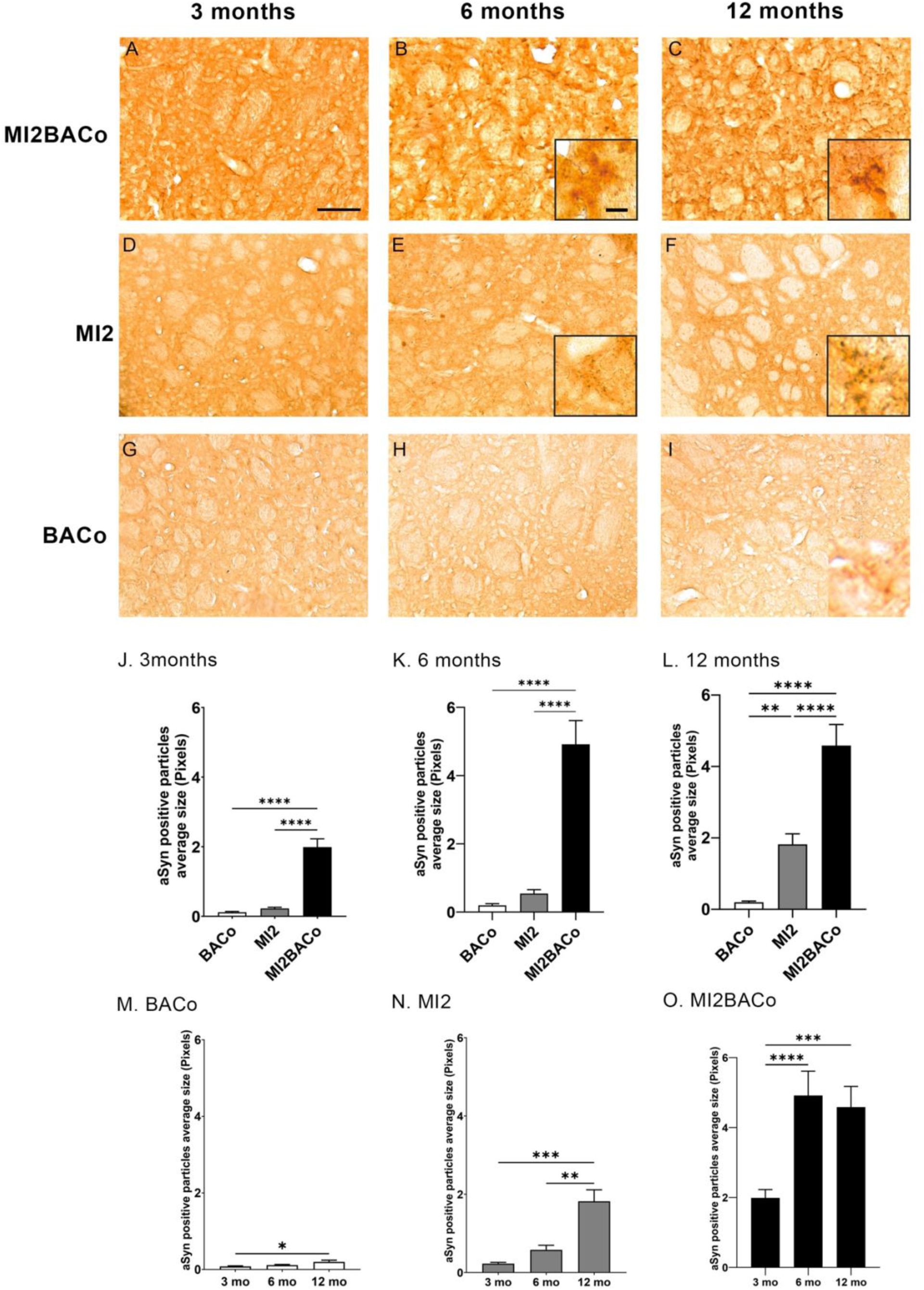
Analysis a-Syn-positive particles size in the striatum of BACo, MI2 and MI2BACo mice (A-I) Representative images showing aSyn immunostaining performed with Syn1 antibody in the striatum of 3-, 6-, and 12-months old MI2BACo (A-C), MI2 (D-F) and BACo (G-I) mice. MI2 and MI2BACo mice exhibit the presence of aSyn-positive clumps, showed in the higher magnifications’ squares (B-C, E-F). Scale bars: 100 μm, 20 μm. (J-L) Graphs are showing results from the analysis of the aSyn-positive particles size in 3 (J), 6 (K), and 12 (L) months old mice. Analysis of particles size shows that, already at 3 months of age (J), MI2BACo exhibited aSyn-positive particles that were increased in size when compared to BACo and MI2 mice. (M-O) Graphs are showing the analysis of the aSyn-positive particles size grouped per genotype, (M) BACo, (N) MI2 and (O) MI2BACo. These graphs show that the dimension of aSyn-positive particles increase with age, also in BACo mice (* p<0.05, ** p<0.01, *** p<0.001, **** p<0.0001, One-way ANOVA+Bonferroni multiple comparisons test. Note different scale in M).

In order to confirm this observation, the striatum of the same mice was stained with the antibody LB509 that specifically recognizes the C-terminus of human aSyn and does not label 1-120haSyn (Figure 3). MI2 mice where not stained by LB509 (Figure 3D-F), while the MI2BACo mice showed the presence of numerous LB509-positive clumps, particularly evident at 6 and 12 months of age (Figure 3, H, I). BACo mice showed some aSyn-staining with the clumps increasing in size overtime even if the amount was significantly lower and size smaller compared to MI2BACo (Figure 3A-C).

**Figure 3.**
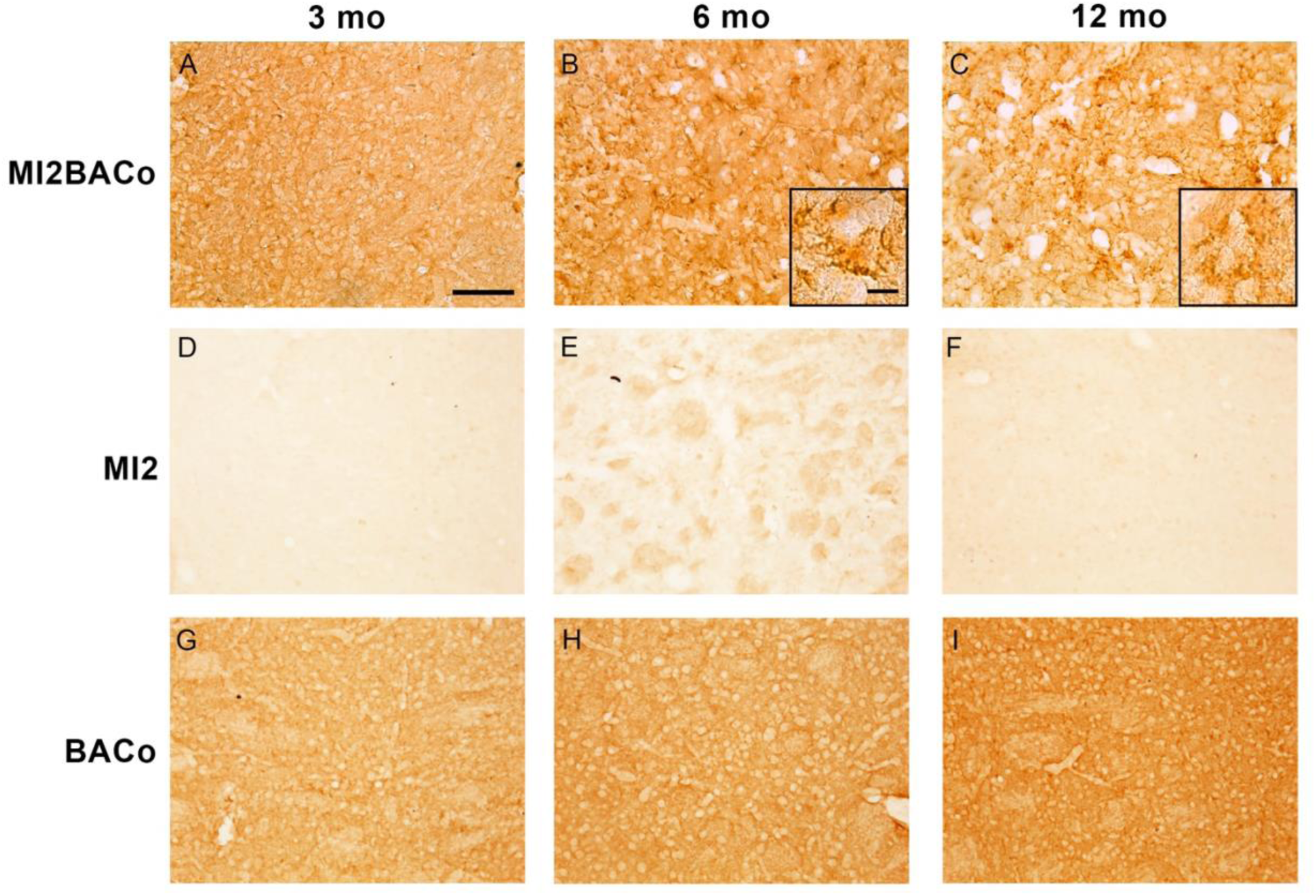
Evaluation of the presence of striatal full-lenght aSyn clumps with LB509 antibody (A-I) Images showing striatal aSyn immunolabeling performed with LB509 antibody, that specifically recognizes full-lenght aSyn (but not 1-120haSyn), in the brain of 3-, 6-and 12- months old MI2BACo (A-C), MI2 (D-F) and BACo (G-I) mice. As MI2 mice expresses only the 1-120haSyn, the striatum of these mice does not display the presence of a specific signal (D-F). BACo mice present a punctate signal, while MI2BACo brains exhibit the presence of LB509-positive clumps more abundant at 6 and 12 months of age (Please see higher magnifications’ squares (B, C). Scale bars: 100 μm in A-I, 20 μm in enlargement (boxed area).

To further evaluate whether the aSyn-positive clumps developed by MI2BACo mice contained fibrillary aSyn, we performed Thio-S staining together with Syn1 and TH immunolabelling (Figure 4). In the striatum of MI2BACo mice we could observe the presence of Thio-S positive signal already at 3 months of age (Figure 4A). Bigger clumps were present at 6 and 12 months of age (Figure 4F, K), and they co-localised with aSyn (Figure 4G, J, L, O). Conversely, BACo mice at 12 months of age did not show consistent Thio-S staining suggesting the absence of fibrillary forms of aSyn, (Figure 4P-T).

**Figure 4.**
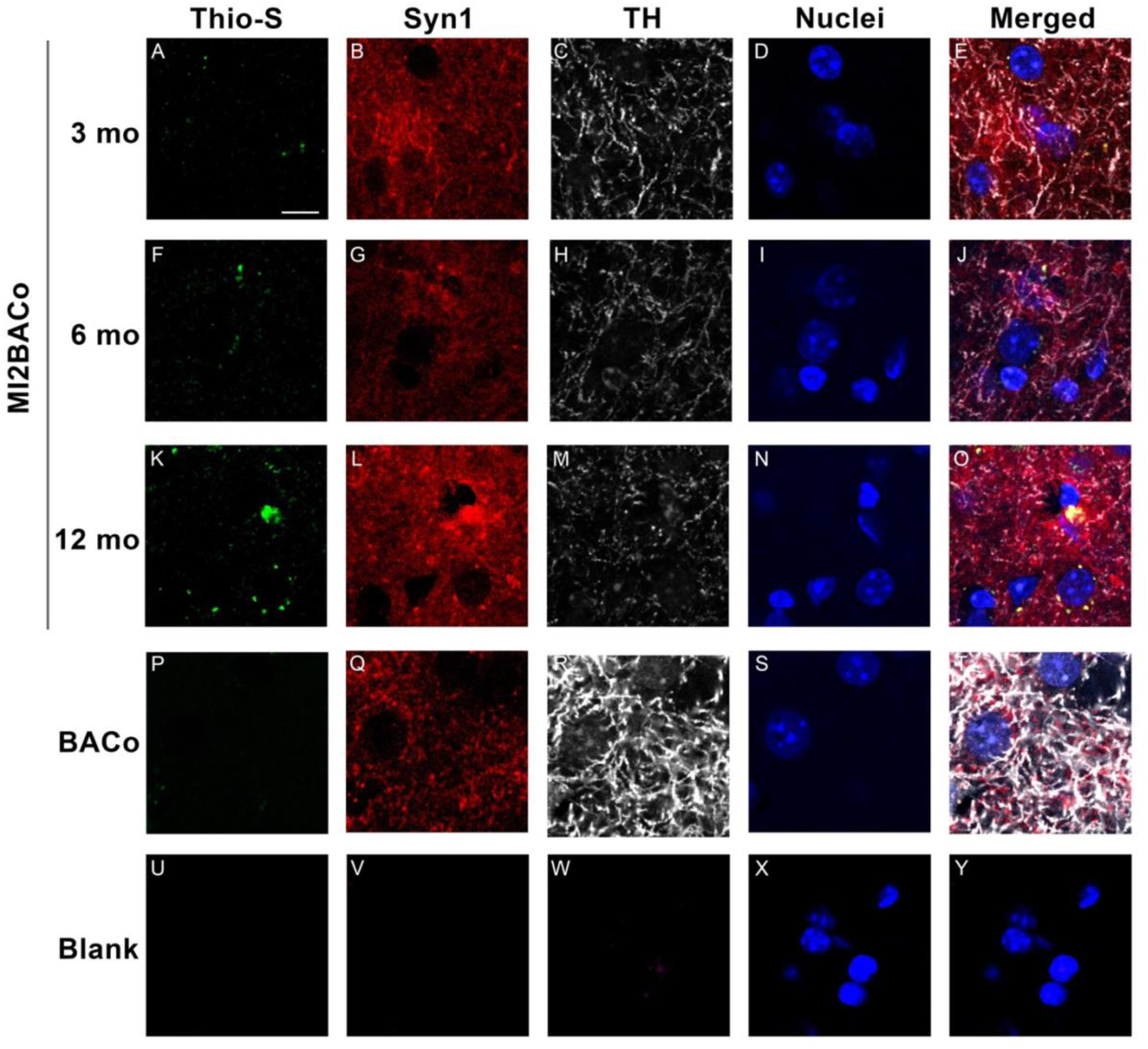
Evaluation of Thio-S-positive staining in the striatum of MI2BACo mice (A-Y) Confocal images showing Thio-S (A, F, K, P, U) staining in Syn1 (B, G, L, Q, V) and TH (C, H, M, R, W) immunolabelled sections from the striatum of 12 month-old MI2BACo and BACo mice. While BACo mice did not reveal the presence of fibrillary/ß-sheet positive aggregates (P), MI2BACo mice exhibited the presence of Thio-S positive staining already at 3 months of age (A). Scale bar: 10 μm.

### The aggregation of aSyn induces a significant redistribution of synaptic proteins in MI2BACo mice

Several studies have shown that aSyn aggregation at the synapse leads to redistribution of SNARE and other presynaptic proteins (Kramer et al. 2007, Garcia-Reitbock et al. 2010, Schulz-Schaeffer 2010, Zaltieri et al. 2015) such as SNAP25 and synapsin III (syn III), a synaptic protein that interacts with aSyn (Zaltieri et al. 2015) and is associated with aSyn fibrils in PD brains (Longhena et al. 2018). We thus investigated whether the co-expression of 120haSyn and full-length aSyn in the MI2BACo mice may enhance syn III accumulation or SNARE protein distribution. We found that all mice exhibited a change in Syn III overtime (Figure 5 J-L). In particular, MI2BACo mice exhibit a marked increase in syn III immunopositive signal from 3 months of age (Figure 5G-I) when compared to BACo (Figure 5 A-C) and MI2 mice (Figure 5D-F). Interestingly we observed that MI2 mice also exhibited an higher accumulation of syn III immunolabelling when compared to BACo mice, especially at 12 months of age.

**Figure 5.**
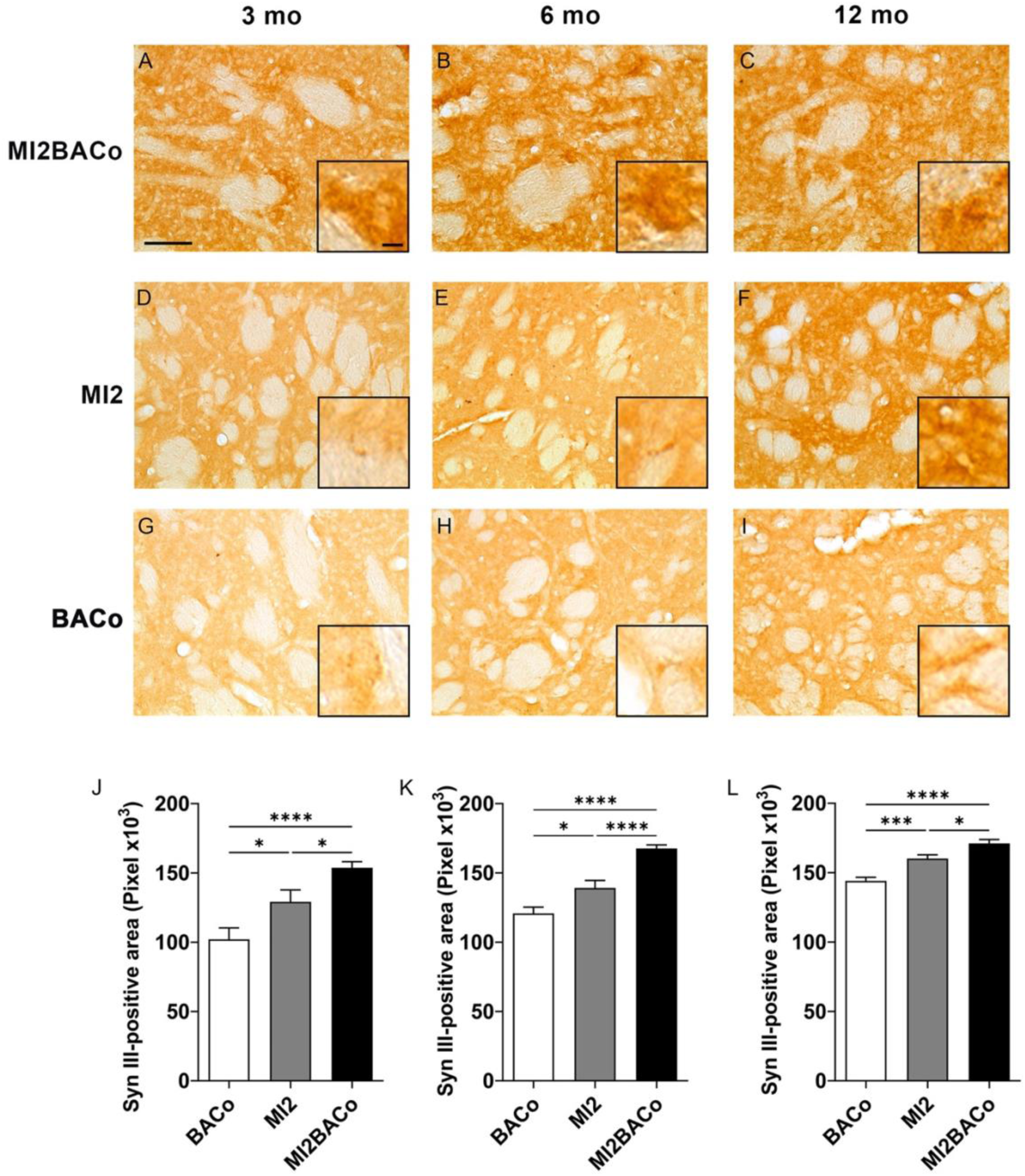
Analysis of striatal syn III in BACo, MI2 and MI2BACo mice (A-I) Representative images of syn III immunostaining in the striatum of 3-, 6-, and 12- months old MI2BACo (A-C), MI2 (D-F) and BACo (G-I) mice. Graphs show results from the quantification of the syn III positive area in 3 (J), 6 (K), and 12 (L) months old mice. Analysis of syn III-positive area shows that both MI2 and MI2BACo mice exhibit increased syn III signal when compared to BACo mice. Results show a significant difference in Syn III staining between MI2 and MI2BACo (* p<0.05, *** p<0.001, **** p<0.0001, One-way ANOVA+Bonferroni multiple comparisons test). Scale bars: 100 μm in A, 20 μm in enlargements.

Study of SNAP-25 distribution in the striatum of BACo, MI2 and MI2BACo mice at 3, 6 and 12 months of age showed the presence of SNAP25-positive clumps in the striatum of MI2 (Figure 6D-F) and in MI2BACo (Figure 6G-I), since 3 months of age. BACo mice only exhibited a few SNAP25-positive clumps at 12 months of age (Figure 6A-C). The results overall indicate a more pronounced aggregation in mouse lines expressing 1-120haSyn.

**Figure 6.**
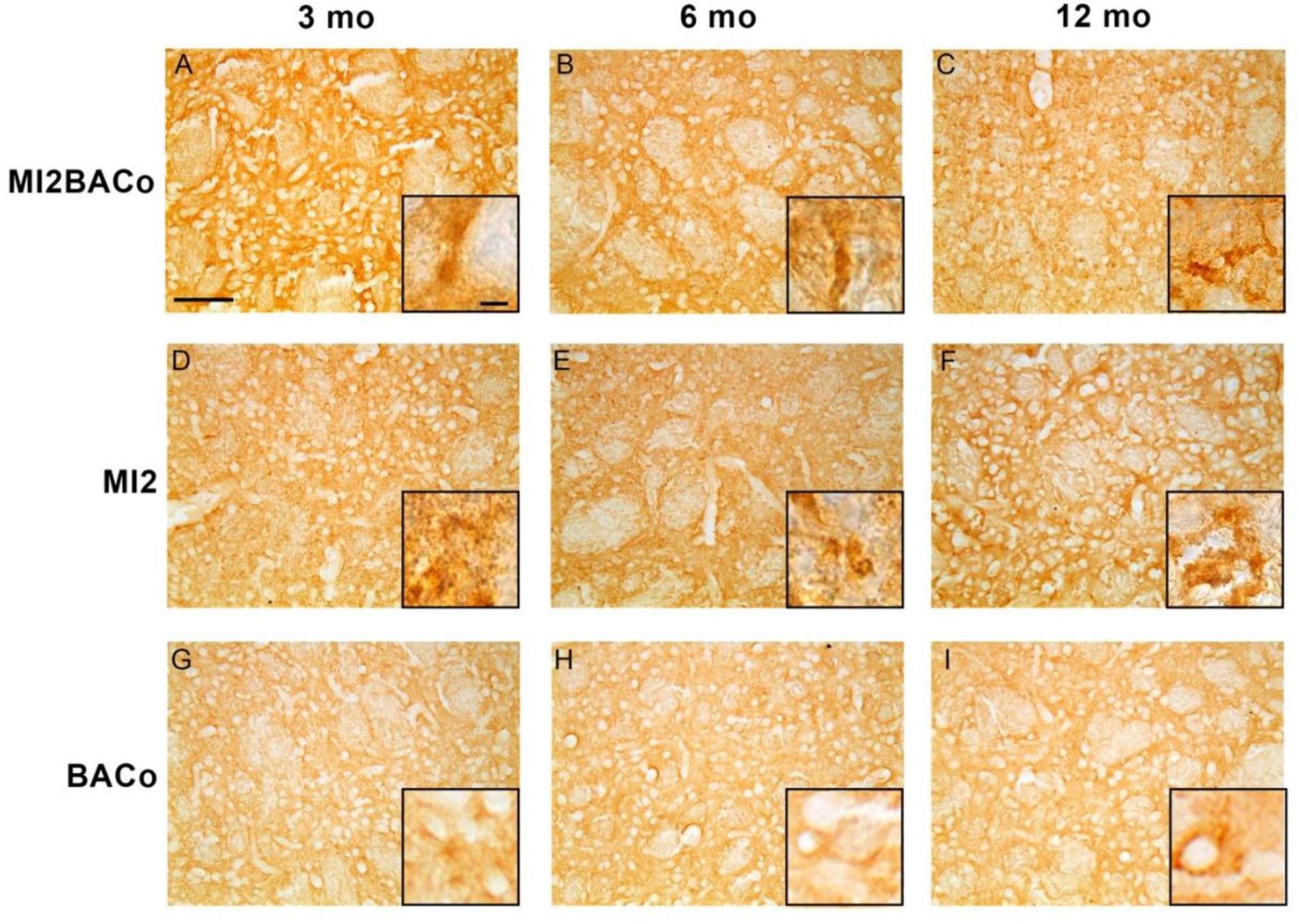
Evaluation of SNAP25 signal in the striatum of BACo, MI2 and MI2BACo mice (A-I) Images showing SNAP25 immunostaining in the striatum of 3-, 6-, and 12-months old MI2BACo (A-C), MI2 (D-F) and BACo (G-I) mice. Labeling in MI2 and MI2BACo brains show the presence of SNAP-25-positive aggregates (Please see boxes with higher magnifications(D-I)). Scale bars: 100 μm in A-I, 20 μm in enlargements.

### Twelve-month old MI2BACo mice show significant loss of TH^+^ dneurons

MI2 mice at 12 months of age show a decreased number of TH^+^ neurons in the *SNpc* and striatal fiber density when compared to the aSyn null OlaHsd mouse line (Wegrzynowicz et al. 2019). To investigated whether the expression of the full-length aSyn influences the effect of 1-120haSyn on dopaminergic neurons we performed TH staining in the nigrostriatal system of BACo and MI2BACo mice at 12 months of age, and the results showed an enhanced nigrostriatal degeneration in MI2BACo compared to BACo mice. Stereological cell count to assess the number of TH^+^neurons in the *SNpc* of MI2BACo, BACo and OlaHsd control mice (Figure 7A-E), showed that MI2BACo mice have a significant decrease of 41% of TH-positive neurons when compared to the control OlaHsd mice (Figure 7D, E). A decrease in TH-positive neurons was also found when comparing MI2BACo and BACo mice but the difference was not significant (Figure 7D, E). A trend to a decrease in TH neurons was also present when comparing BACo with OlaHsd although the difference was not significant (Figure 7 D, E).

**Figure 7.**
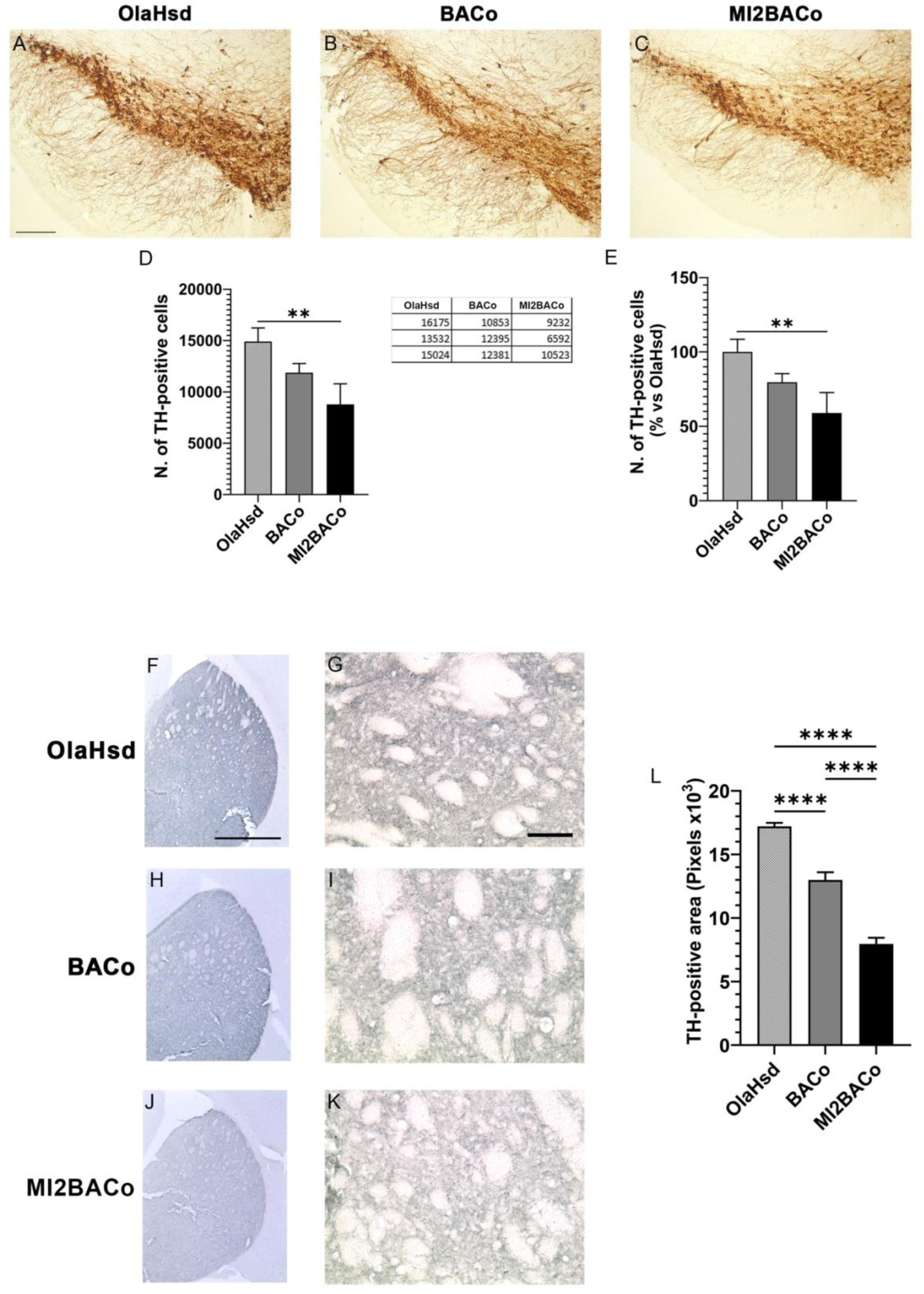
Analysis of nigral TH^+^ neurons and striatal fibers (A-C) Representative images of TH immunolabeling of the *substantia nigra* of 12-months old Ola, BACo and MI2BACo mice. Scale bar: 500 μm. (D, E) Graphs showing the analysis of TH+ neurons number in the *substantia nigra pars compacta*. The results reveal that MI2BACo mice show a significant decrease in TH+ neurons number when compared with OlaHsd mice (**p<0.01, One-way ANOVA+Bonferroni multiple comparisons test). (F-K) Striatal TH staining in the brain of 12-months old OlaHsd (F, G), BACo (H, I) and MI2BACo (J, K) mice. Scale bars: 500 μm (F, H, J), 20 μm (G, I, K). (L) Graph presenting the analysis of TH^+^ striatal area. MI2BACo mice show a significant reduction in TH^+^ area when compared with BACo and OlaHsd mice. Of note, BACo mice also exhibit a significant TH^+^ decrease in TH^+^ processes when compared with OlaHsd control mice (**** p<0.0001, One-way ANOVA+Bonferroni multiple comparisons test).

We then analyzed dopaminergic arborization in the striatum using TH immunolabeling (Figure 7F-L). The results showed a significant reduction of TH^+^ fibers in the striatum of 12month-old MI2BACo mice when compared to both OlaHsd and BACo mice, Interestingly, in line with the evidence supporting that aSyn aggregation induces a retrograde degeneration pattern, we found that BACo mice also exhibit a significant decrease of TH^+^ fibers when compared to OlaHsd mice.

## DISCUSSION

Transgenic mouse models are commonly used to investigate mechanisms of disease in PD and other neurodegenerative disorders (Rocha et al. 2023). Here we describe two new transgenic mouse models, that express human full-length aSyn under the human aSyn promoter on a null aSyn background. While BACo mice express only human full-length aSyn, MI2BACo mice, also express the human 1-120haSyn under the rat TH promoter, resulting from the crossing of the BACo mice and the previously described MI2 mice (Wegrzynowicz et al. 2019).

The main aim of this study was to determine if the presence of human full-length and C-terminally truncated aSyn may enhance human full-length a Syn aggregation and toxicity *in vivo*. Initially we looked at the presence of aSyn aggregates in the striatum of BACo and MI2BACo mice compared to MI2 mice using immunohistochemistry. Because both the 1-120 truncated and full-length aSyn are human, we needed an antibody that could differentiate between them, and we raised the L91 antibody that specifically recognizes the 1-120haSyn.

We found that MI2BACo mice had more aSyn aggregates compared to both MI2 and BACo mice. Some of the aggregates were clearly labelled by L91 antibody revealing the presence of the truncated protein. Moreover, aggregates in the MI2BACo appeared earlier, being present at 3 months of age and increasing overtime, with some aggregates in the striatum becoming Thio-S positive. The aggregates in the MI2 mice at 12 months of age were comparable in size to those of MI2BACo mice at 3 months of age. BACo mice showed a significant increase in the aggregates between 3 and 12 months of age, but their sizes were significantly reduced compared to the MI2 mice at 12 months of age. When compared to MI2BACo mice, aggregates in BACo mice were reduced at all ages, suggesting a decreased aggregation of full-length human aSyn compared to the 1-120 truncated protein. In MI2BACo mice the redistributions of synaptic proteins, such as Syn III and SNAP25, previously reported in the 1-120haSyn transgenic mice (Garcia-Reitbock et al. 2010; Wegrzynowicz et al. 2019; Faustini et al. 2020) were also significantly increased.

In PD aSyn aggregation is believed to start in the synapses in the striatum and to progress retrogradely to the cell bodies of dopaminergic neurons in the SNpc resulting in their dysfunction and death. We therefore looked at the amount of TH^+^ processes and terminals in the striatum in BACo and MI2BACo mice and found that they were significantly reduced in MI2BACo compared to controls and BACo mice. A reduction was also present in the BACo mice compared to control mice but was less pronounced than that present in the MI2BACo mice.

When we counted the number of TH^+^ neurons in the *SNpc*, we found that they were significantly decreased in MI2BACo mice compared to OlaHsd controls but not compared to BACo mice. BACo mice showed a trend towards a decrease of TH^+^ neurons which, however, did not reach significance when compared to control mice. Previously we reported a 31% loss of dopaminergic neurons in *SNpc* of MI2 mice (Wegrzynowicz et al. 2019), that is milder than that found in MI2BACo mice, indicating that the presence of full-length aSyn enhances the dopaminergic neuronal death induced by 1-120haSyn. These results suggest that in the BACo mice the ongoing pathological process - is less advanced than in MI2 and MI2BACo mice. In a different BAC mouse model, the SNCA-OVX mice, similar to the BACo mice in that they express similar amounts of aSyn in a null aSyn background, 30% of cell loss was present at 18 months of age (Jazenic et al. 2013). It will be interesting to investigate TH^+^ neuron numbers in the *SNpc* of BACo mice at a similar age to determine whether a significant loss is also present.

By contrast, in another BAC mouse expressing the same transgene as the one used in the BACo mice, but with mouse aSyn in the background, no TH^+^ neuron loss in the *SNpc* was observed at 24 months of age (Yamakada et al. 2012). It is possible that the presence or absence of mouse aSyn in the background can influence the development of pathology and cell loss. Indeed, it has been reported that mouse aSyn inhibits the aggregation of human aSyn Luk et al. 2016; Ohgita et al. 2023; Sokratian et al. 2024). However, it could be also possible that the initial aggregation of human aSyn in a transgenic mouse lacking endogenous aSyn creates an initial toxic loss of function due to the reduction of the protein which is then aggravated by the toxic gain of function of the aggregates.

Interesting, the presence of the mouse aSyn is important during development in that its absence causes a small but significant reduction of the number of dopaminergic neurons in the SNpc (Garcia-Reitbock et al. 2013). Re-expressing mouse aSyn in SNCA KO mice rescues this reduction in a dose-dependent manner (Garcia-Reitbock et al. 2013). No increase of TH^+^ neurons is found in the BACo mice that are in a null mouse aSyn background, suggesting that the presence of full-length human or mouse aSyn, do not have the same effect in a mouse.

In summary, the MI2BACo mice show faster and significantly higher protein aggregation, loss of TH^+^ striatal processes and dopaminergic neuron death in the *SNpc* compared to BACo, indicating that the 1-120 truncated and aggregation-prone aSyn may play a role in the pathological process also in the human brain.

The results shown indicate that the MI2BACo mice recapitulate the major pathological features of PD and are a good model to study both mechanisms of disease and to test treatments targeting aSyn pathology.

## Acknowledgements

This work was supported by a grant from Parkinson’s UK (grant G-1703) and the Scholl Foundation. We are grateful to Professors Michel Goedert and Aviva Tolkovsky for helpful comments. Part of the confocal imaging experiments were performed at the Imaging Platform of the Department of Translational and Molecular Medicine of the University of Brescia (Italy).

## SUPPLEMENTARY FIGURES

**Supplementary Figure 1.**
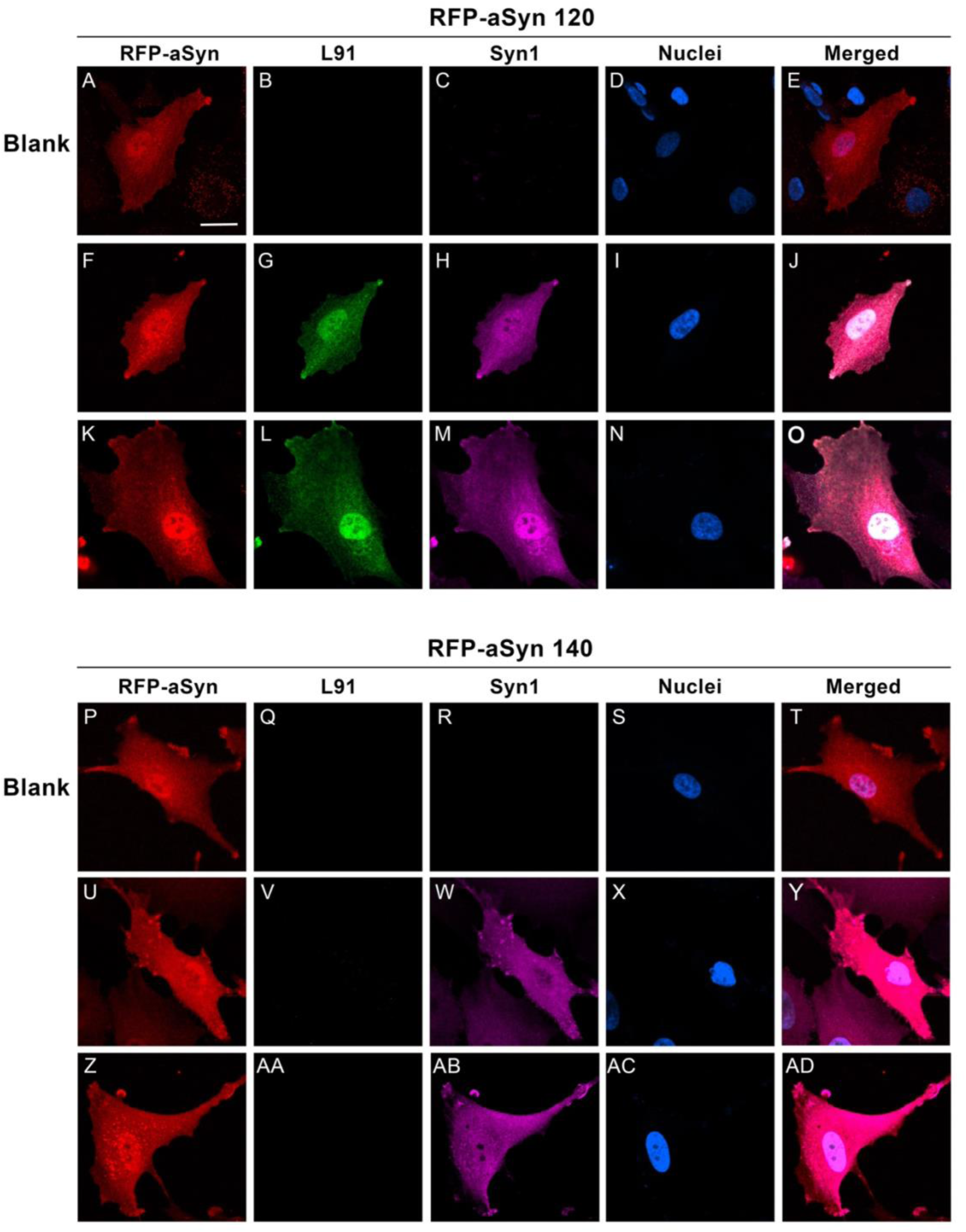
Staining of L91 and Syn1 in SK-N-SH cells overexpressing full length or 1-120 truncated aSyn. (A-AD) Images showing L91 immunolabeling of SK-N-SH cells overexpressing truncated 1- 120 aSyn tagged with RFP (SK aSyn 120, A-O) or full-length aSyn tagged with RFP (SK aSyn 140, P-AD). The staining confirms that L91 specifically recognises truncated 1-120 human aSyn. Scale bar: 10 μm.

**Supplementary Figure 2.**
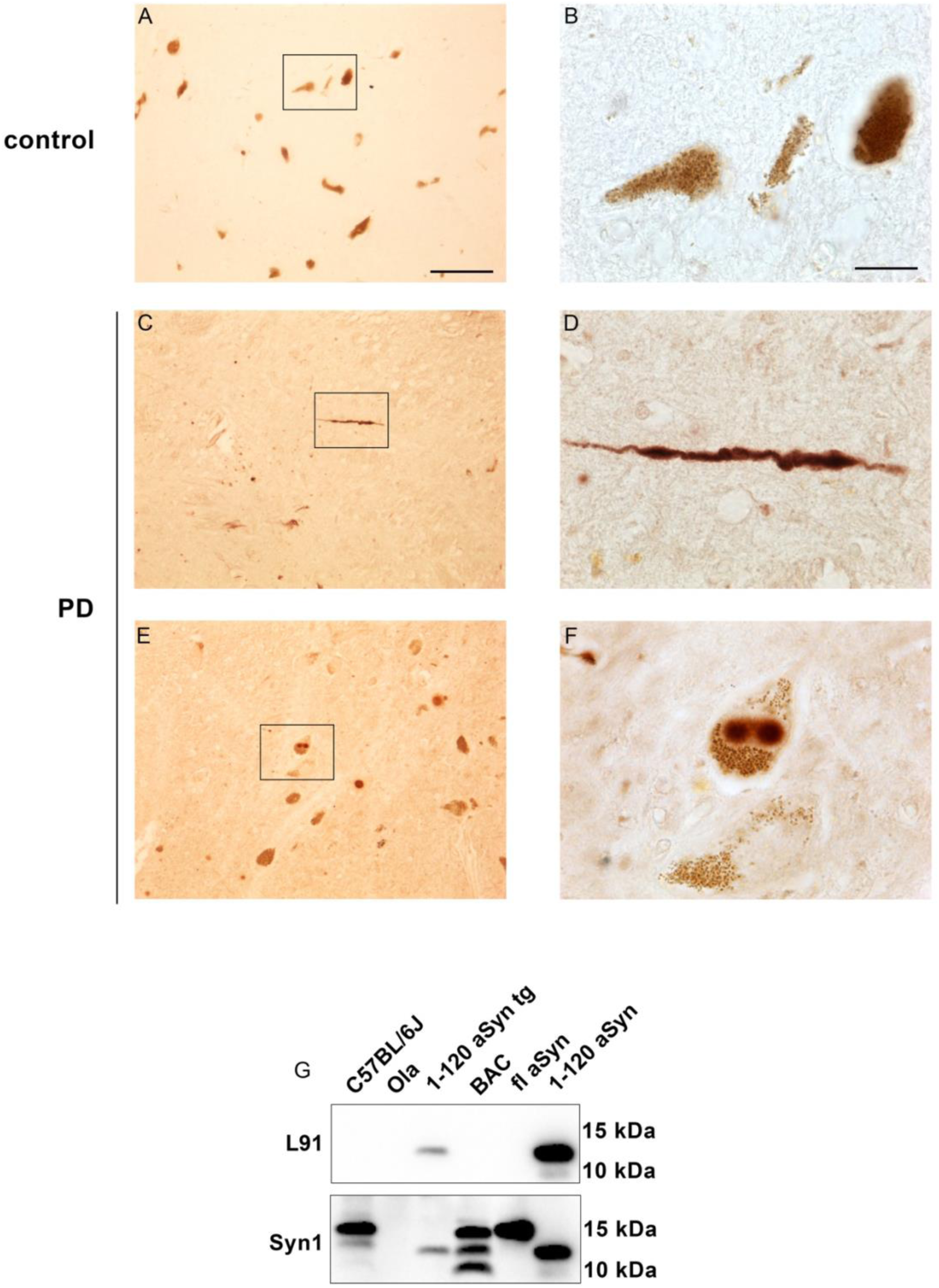
L91 staining in PD and control human brains. (A-F) Images show L91 immunolabeling in post-mortem substantia nigra of two patients affected by PD (C-F) and one age matched control (A, B). Truncated aSyn-specific antibody shows the presence of LN (C, D) and LB (E, F) in the brain of PD patients. Scale bars: 100 μm in (A,C,E), 20 μm (B,D,F). (G) Representative L91 and Syn1 immunoblots performed on nigrostriatal protein extracts of 12 months old C57Bl6J, BACo, 1-120 aSyn Tg,OlaHsd mice and recombinant full-length and truncated 1-120haSyn. The L91 polyclonal antisera recognizes human 1-120-haSyn in transgenic mice and the 1-120 recombinant protein.

**Supplementary Figure 3.**
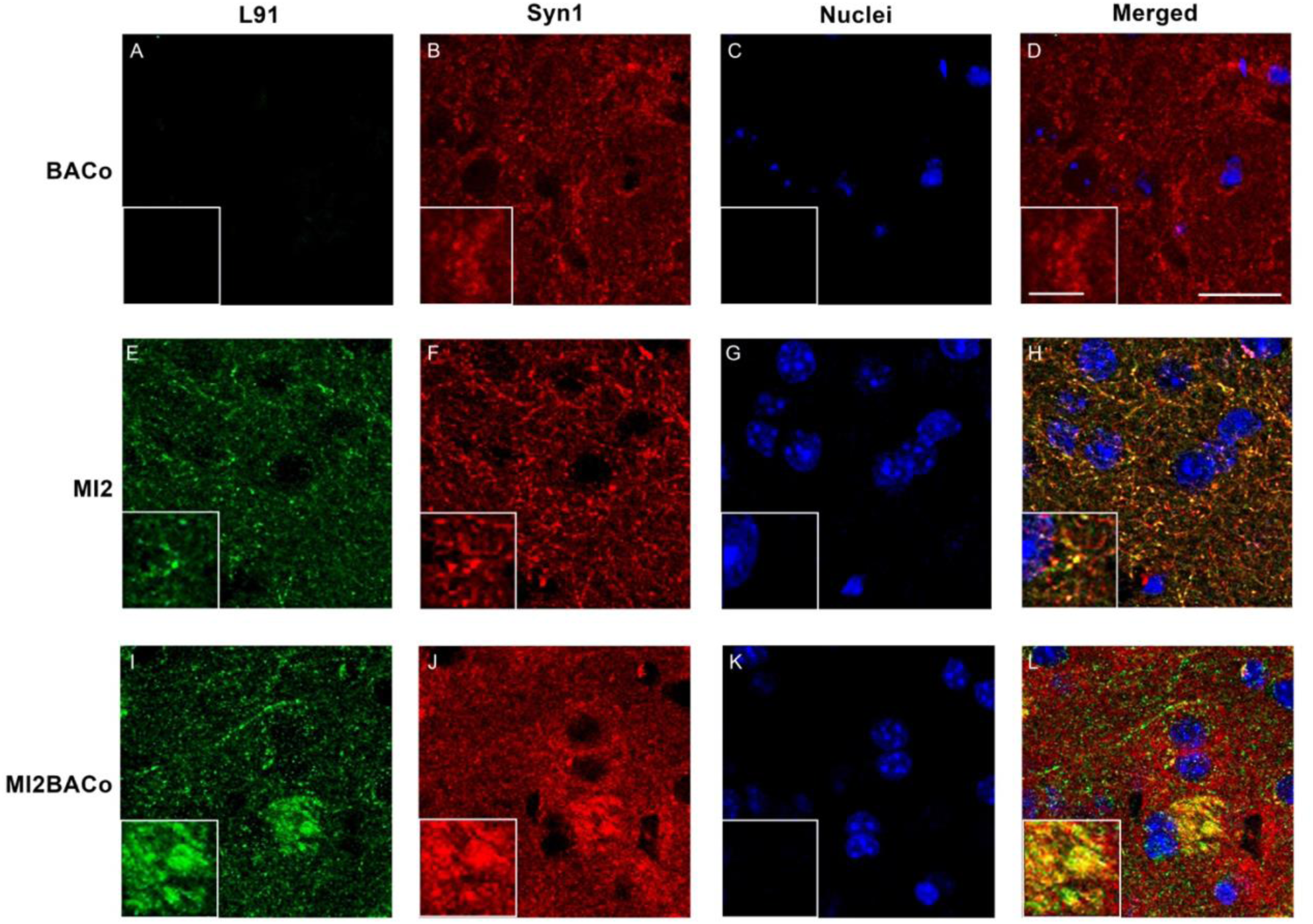
Co-localization of L91 and syn1 antibody inn BACo, MI2 and MI2BACo mice (A-L) Representative confocal images showing co-labelling with L91 and Syn1 of striatal sections of 12 months old BACo, MI2 and MI2BACo mice. In MI2 and MI2BACo mice, L91 signal colocalises with aSyn staining showing the presence of truncated 1-120 aSyn-positive aggregates (Higher magnifications’ squares, E-H, I-L). Scale bars: 20 µm in A-L and 5 μm in enlargements.

**Supplementary Figure 4.**
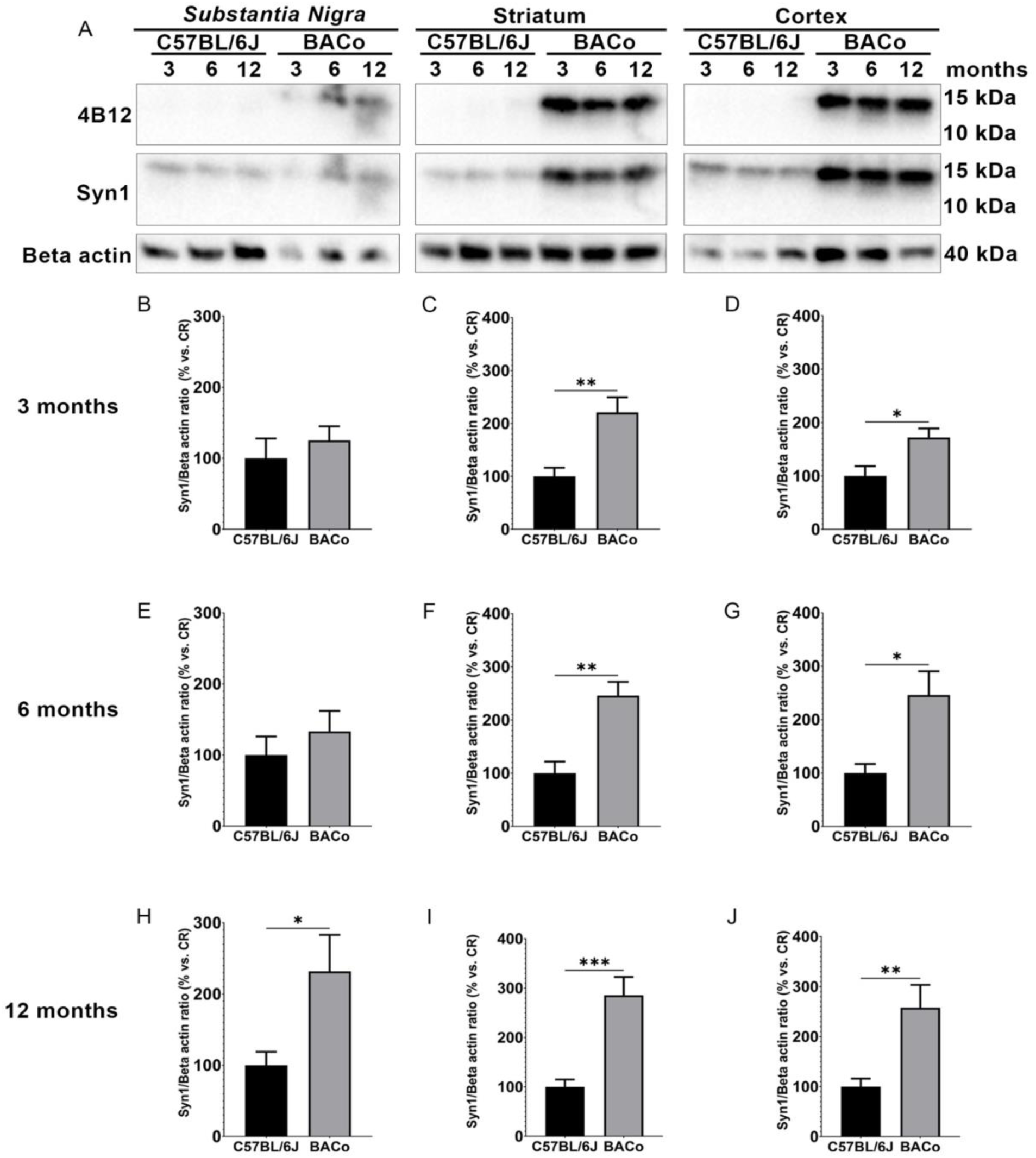
Expression of human alpha-synuclein in brain of BACo mice compared to C57Bl6J (A) Representative immunoblots of 4B12, an antibody that specifically recognizes human aSyn, or Syn1 that recognizes both truncated and full-length human and mouse aSyn and beta-actin performed on protein extracts from *substantia nigra*, striatum and cortex of 3-, 6- and 12-months old C57Bl6J wild type and BACo mice. (B-J). Graphs show Syn1 protein levels normalized to beta actin in 3 (B-D) 6 (E-G) and 12 (H-J) months old mice substantia nigra, striatum and cortex. Results reveal a 2-to-3-fold increase of aSyn levels in BACo mice when compared to CR mice. This increase is significant in the *substantia nigra* of 12-months old mice (H), in the striatum (C, F, I) and in the cortex (D, G, J) (* p<0.05, ** p<0.01, Unpaired t-student test, 3 mice per group).

